# The interrelation between the 3D microenvironment and mechanics of human induced pluripotent endothelial progenitors

**DOI:** 10.1101/2025.06.18.660409

**Authors:** Toni M. West, Jiwan Han, Gabriel Peery, Janet Zoldan, Michael S. Sacks

## Abstract

Human induced pluripotent stem cells (hiPSCs) offer immense potential for tissue engineering, yet poor vascularization remains a significant hurdle. Understanding how hiPSC-derived endothelial progenitor cells (hiPSC-EPs) form networks is essential for therapeutic progress. This study investigates extracellular matrix (ECM) remodeling and cellular contractility during the early self-assembly of hiPSC-EPs within 3D hyaluronic acid-based hydrogels. By tracking microsphere displacements before and after cytochalasin-D treatment, we quantified contractile forces in single cells and clusters at days 4 and 7. We then applied a novel inverse modeling approach using a compressible material model to determine spatially varying changes in hydrogel modulus caused by enzymatic degradation and ECM deposition. Our findings reveal that basal contractility and remodeling are nonlinearly influenced by multicellularity, culture duration, and initial stiffness. Increased hydrogel stiffness, paired with synergistic rises in strain energy, resulted in high traction forces in longer culture times. These results provide critical mechanical insights into hiPSC-EP self-assembly, advancing our ability to engineer functional vascular networks.

## 1. Introduction

Human induced pluripotent stem cells (hiPSCs) offer promising untapped patient-specific and immune-evasive cell sources. Therefore, hiPSCs are emerging key players in regenerative medicine and research, not only because they hold the potential to produce a variety of tissue types, but also because of their potential to be engineered for immune compatibility and their projected capacity for large-scale production. [1, 2] However, one of the major challenges in clinical translation of hiPSC-based tissue therapies is the lack of vascularization within the derived tissues. This lack of microvasculature in hiPSC-derived constructs has even greater ramification when attempting to repair or study highly vascularized systems that also contain tissue-specific endothelial cells such as heart or brain tissue. [3, 4] Therefore, developing approaches to induce vascularization will not only improve the efficacy of hiPSC-derived tissues, but it could also become a therapy in and of itself for primarily vascular-related disorders including ischemia, embolism, and diabetic vasculopathy. These challenges underscore the urgent need to deepen understanding of the process of vasculogenesis in hiPSC-derived cells.

Endothelial cells form the capillaries of the vascular system. Methods developed by our group and others differentiate hiPSCs into endothelial progenitor (EP) or endothelial cells (ECs), producing cells with varying maturation and tissue-specific properties depending on the protocol. [5, 6] When hiPSC-EPs are embedded in hyaluronic acid (HA)-based hydrogels and supplemented with vascular endothelial growth factor (VEGF), they undergo morphogenesis into interconnected spindle-shaped structures that form lumens through extracellular matrix (ECM) remodeling. [1, 7, 8] In norbornene-functionalized HA (NorHA) hydrogels cross-linked with enzyme-degradable peptides (EDPs), we previously found that more compliant hydrogels (190 Pa) promote greater vascular connectivity, longer branches, and enhanced actin remodeling, whereas stiffer hydrogels (551 Pa) increase mechanotransduction, as indicated by YAP/TAZ nuclear localization and elevated CD31 expression. [9] These results demonstrate that hydrogel stiffness regulates vasculogenesis as hiPSC-EPs mature into *CD*31^+^ endothelial cells.

However, these studies did not consider how cells remodel their microenvironment. We and others have shown that cells can significantly modify the local modulus of 3D hydrogels [10–13] and actively remodel the extracellular matrix through deposition of proteins such as collagen and laminin, as well as secretion of degradation enzymes including matrix metalloproteinases (MMPs) [5, 14–16]. Consequently, hiPSC-EPs are expected to locally alter the mechanical properties of NorHA hydrogels during culture. Importantly, these mechanically driven processes emerge not only at the single-cell level but also through collective multicellular interactions that govern tissue-scale organization. How hydrogel stiffness, culture duration, and multicellularity together regulate the transition from single-cell mechanosensing to coordinated network-level mechanical responses that enable microvascular assembly by hiPSC-EPs remains largely unknown.

As a first step toward addressing these questions, this study characterized ECM remodeling and basal contractility of hiPSC-EPs in EDP-crosslinked NorHA hydrogels of varying stiffness across different culture durations and cell densities. Building on our previous methods for quantifying cell contractile behavior [17], we measured basal contractile displacements and hydrogel kinematics induced by hiPSC-EPs. We also applied recently developed approaches for determining spatial variations in hydrogel modulus [18, 19] to more accurately calculate strain energy density and traction forces. We hypothesized that, prior to vascular network formation, multicellular cooperativity would increase hiPSC-EP basal contractility over time and that these effects would depend on the initial shear storage modulus of the hydrogel. We further expected that multicellular systems would exhibit synergistic increases in ECM remodeling, actin cytoskeleton maturation, strain energy density, and traction forces. Together, these analyses advance understanding of the mechanical cues that regulate microvascular development.

## 2. Methods

In this study, we quantified contractile and ECM remodeling behaviors of hiPSC-EPs in 3D hydrogels during early capillary-like assembly using 3D traction force microscopy (Figure 1). tdTomato-expressing hiPSCs were differentiated into EPs [5, 20] and embedded in NorHA hydrogels with fluorescent microbeads, cross-linked with enzyme-degradable peptides at varying percentages to produce different stiffnesses [9]. Cells were imaged at 4 or 7 days post-embedding, and FM-TRACK [17] tracked marker displacements from basal contractility cessation to compute local 3D hydrogel deformations. Cell surface tractions were estimated via an inverse computational model. Unlike traditional TFM, which assumes homogeneous hydrogel mechanics [21–23], we accounted for hydrogel remodeling by enzymatic degradation and ECM deposition [18, 19], applying novel inverse modeling to resolve spatially varying stiffness and accurately quantify cell tractions.

**Figure 1.**
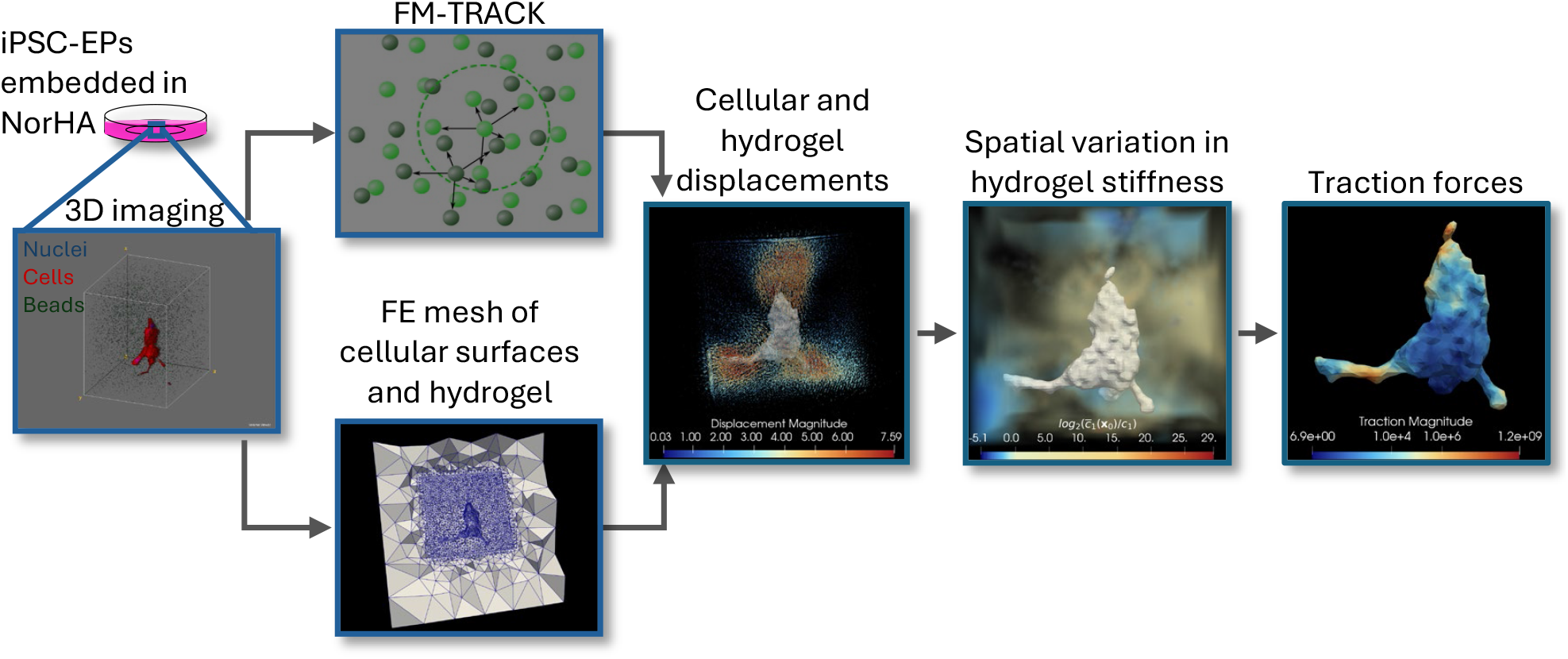
Pipeline for Experimentation and Analysis. Schematic overview of the experimental and analytical workflow implemented to produce the presented data. Induced pluripotent stem cell-derived endothelial progenitor cells (hiPSC-EPs) were embedded in norbornene-functionalized hyaluronic acid hydrogels (NorHA) containing green fluorescent beads which were cultured for 4 or 7 days. Samples were imaged via 3D confocal microscopy under both basal and cytochalasin-D treated states. FM-TRACK software [17] analyzed the position and displacement of the beads in the hydrogel. The confocal’s blue channel was evaluated to quantify cell number, and the red channel was utilized to produce a finite element (FE) mesh of the cells and surrounding gel. Displacements were then mapped onto the nodes of the cell and hydrogel meshes, and a kinematic analysis of the hydrogel was completed. 190 Pa hydrogel systems were then further analyzed to determine local hydrogel modulus, system strain energy, and cellular traction forces.

### (a) Differentiation and Gelation of 3D Endothelial Progenitor Cells (EPs)

hiPSCs stably expressing tdTomato (WiCell) were differentiated into EPs as previously described by Jalilian et al. [20] and us [5]. On day 5 of differentiation, cells were sorted for EP marker, CD34. Sorted cells were embedded in NorHA hydrogels containing dragon green 0.51 *µ*m diameter beads (Bangs Labs) as we have described in detail [9]. Three types of hydrogels were produced with varying concentrations of the enzymatically degradable peptide (EDP) KCGPQGIWGCK, with 25%, 50%, and 75% cross-linked hydrogels having stiffnesses of 190 ± 33.3 Pa, 336 ± 67.4 Pa, and 551 ± 90.1 Pa, respectively [9]. Cell-laden hydrogels were cultured in Endothelial Cell Growth Medium (EGM-2, Promo Cell) supplemented with 50 ng/mL vascular endothelial growth factor (VEGF165; Acro Biosystems) and 10 *µ*M ROCK inhibitor Y-27632 (Selleckchem) for the first 24 hours and were changed daily thereafter with EGM-2 media for either 4 or 7 days.

### (b) 3D Traction Force Microscopy (3D-TFM)

3D-TFM was performed on single cells and small multi-cell groups that fit within the microscope field of view (FOV) from hiPSC-EP-laden hydrogels cultured for either 4 or 7 days, as we have previously described [24] with the following improvements. Immediately prior to imaging, the embedded hiPSC-EPs were stained for 30 minutes with Hoecht following manufacturer directions. Then, 2.9 mL of phenol-free media was placed in the dish. Images were taken with a 40x silicone oil objective lens on a Nikon AXR confocal equipped with a Tokai hit stage incubator set to 37°C and a Queensgate NanoScan SP Z Series Piezo. A 147×147×119.23 *µ*m FOV with isotropic voxels of side length 0.288 *µ*m that had been corrected for spherical aberration was imaged. Single cells or multi-cell groups were identified by the number of nuclei that were associated with the tdTomato-expressing portions of the image. After hiPSC-EPs were imaged in their basal states, the potent actin polymerization inhibitor, cytochalasin-D (CytoD, MilliporeSigma) was added to achieve a 4 *µ*M concentration. After 40 minutes of CytoD treatment at 37 °C, the same FOV was then re-imaged with the cells in the fully-relaxed steady state.

### (c) Displacement Analysis

Confocal images were processed with the FM-TRACK software we have previously described, [17] which tracks the movement of the fiducial marker beads embedded in the hydrogel going from the fully-relaxed, CytoD-treated steady state to the basal state so that basal contractility can be assessed. A Gaussian process regression (GPR) model that accounted for the inherent noise of the microscope [25] was produced of the cell/hydrogel location-dependent displacements. Displacements were then interpolated onto the cell surface FE mesh to establish a direct one-to-one correspondence between the fully-relaxed steady state of cells and their basally-contracted state. To assess the direction of contractile displacement relative to the surface of the cell, signed displacement magnitudes were computed by multiplying displacement magnitudes by the dot product of unit vectors pointing radially outward from the cell center with nodal cellular displacement vectors. To graph contractile response, the normalized displacements were multiplied by -1 to make contractile events positive, and then a threshold of 0 was set to isolate contractile regions.

For analyzing the kinematic parameters of the NorHA hydrogels, a finite element (FE) mesh of each hydrogel was generated with Gmsh python library functions. [26] A multi-tiered bounding box approach was employed to produce the hydrogel mesh, producing a computationally tractable model with uniform dimensions. The inner-tier box represented the experimentally-imaged FOV around the cell(s), while the outer-tier box represented extrapolated readouts of what occurs in the hydrogel 150 *µ*m from the cell(s) (Figure 1). The element side lengths from the cell surface to the edge of the inner bounding box were set to average 2.86 *µ*m. This characteristic side length was determined from an exemplar cell to balance geometric fidelity and computational efficiency. Implementing similar methods, the element side lengths between the inner and outer bounding boxes were set to increase exponentially from 2.86 *µ*m to 50 *µ*m. The cell itself was modeled as a hollow inclusion in the center of the hydrogel. The GPR model was then evaluated at each node of the hydrogel FE mesh. The directions of signed displacement vectors were normalized to the cell centroid as described above to ensure consistent directional interpretation. The resulting normalized displacements were then parsed by standard deviation of displacement magnitudes throughout the hydrogel to evaluate the spatial distribution of contractile vs. expansile regions of the cell.

### (d) Hydrogel Deformation Analysis

Because the hydrogel may have properties of a compressible solid, an assessment of the deformation behavior of the NorHA hydrogel was performed by calculating the Jacobian determinant, which quantifies local volumetric changes. The deformation gradient was given by **F** = **I** + ∇**u** for each element in a given hydrogel mesh, where **I** is the identity tensor and ∇**u** is the displacement gradient tensor. This was computed in Python according to the constant strain tetrahedron approach, multiplying the matrix of relative displacements between the nodes of a tetrahedron by the inverse of the matrix of relative reference positions. The right Cauchy-Green deformation tensor was then calculated as **C** = **F**^*T*^ **F**, and the Jacobian determinant was defined as *J* = det(**F**).

### (e) Kinematic Analyses

To determine the extent to which volumetric strain plays a role in these iPSC-EP/NorHA hydrogel systems, a decomposition of the volumetric and deviatoric portions of strain was performed. To accomplish this, standard invariant measures of large deformation, such as the first invariant of the right Cauchy-Green deformation tensor, *I*_1_ = *tr*(**C**), where **C** = **F**^*T*^ **F**, were modified to account for *J* ≠ 1. Thus, for compressible hydrogels *I*_1_ was adjusted to account for the observed volume changes

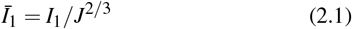

From this relationship, we computed a total deviatoric strain index for the entire hydrogel volume using

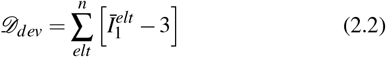

to quantify the shape-distorting component of the deformation independently of volume effects. Similarly, to account for the total changes in hydrogel volume from cellular contraction, the total volumetric strain index for the entire hydrogel volume was defined as

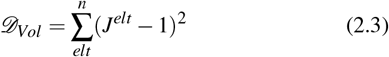

### (f) Inverse Modeling of Spatially Varying Hydrogel Stiffness

Spatial variations in cell-modified hydrogel stiffness were then determined so that strain energies and traction forces could be accurately calculated for the hydrogels and iPSC-EPs, respectively. The spatial variation in stiffness, once calculated, was also utilized as a source for understanding the ECM remodeling potential of the iPSC-EPs as they begin to form into capillary networks. Inverse modeling approaches that we have previously described [18, 19] were implemented to determine the changes in stiffness in the hydrogels. In brief, the GPR model described above of displacements was evaluated at all points in a mesh consisting of only the imaged region of hydrogel, with spline lengths averaging 2.86 *µ*m. Displacement magnitudes of *>* 0.38*µ*m within each hydrogel served as the ground truth values for the inverse model to match since lesser values have significant inherent noise from the imaging system associated with them. The region of hydrogel with *>* 0.38*µ*m was defined as within the “event horizon.” Because the event horizon volumes needed to be a substantial proportion of the hydrogel volumes for the model to produce accurate results, only the 190 Pa samples were analyzed. In the forward-solves performed in FEniCS 2019.1.0, the hydrogels were modeled as neo-Hookean hyperelastic compressible solids, which varied spatially according to location in the hydrogel, **x**_**0**_, such that

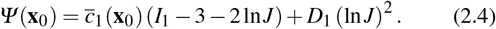

*D*_1_*/c*_1_ = 1 was implemented to match the observed compressibility of the hydrogels. The value of the material constant, *c*_1_, corresponded to half the experimentally measured shear modulus, consistent with the Neo-Hookean small-strain relation, *µ* = 2*c*_1_. Furthermore, to improve numerical stability over several orders of magnitude, 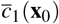, was given exponential form,

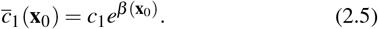

The field *β* (**x**_0_) was discretized on the hydrogel mesh using linear Lagrange basis functions, with values stored at each node.

The inverse analysis employed the limited-memory Broyden-Fletcher-Goldfarb-Shanno (L-BFGS) algorithm [27]. Starting from *β* (**x**_0_) = 0, the algorithm updated *β*_*sim*_ to minimize the

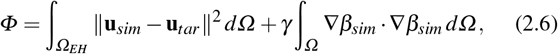

where *Ω* denotes the imaged domain and *Ω*_*EH*_ denotes that within the event horizon. Gradients of *Φ* for L-BFGS were generated with FEniCS-adjoint [28] and MOOLA [29], with spatial gradient weighting applied to maintain numerical stability and promote smooth variations in *β* (**x**_**0**_). Pilot tests indicated that this additional smoothing was necessary beyond the Tikhonov term in Eq.(2.6). Displacements at each iteration were obtained by minimizing the strain-energy functional

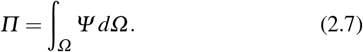

Experimentally measured boundary displacements were imposed as Dirichlet conditions on the MVIC surface and hydrogel boundary, ensuring agreement with target values. Convergence thresholds were selected based on prior ground-truth studies, where only 1.6% of nodes exceeded displacement errors of 0.38*µ*m and each hydrogel required approximately 1 h of computation [19]. Once the optimization returned the 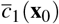 field, the *log*_2_ fold change of stiffness was then calculated as 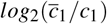.

### (g) Statistical Analysis

A value of *p* < 0.05 was considered significant. Implemented statistical tests are delineated in each figure legend.

## 3. Results

### (a) hiPSC-EP Basal Contractile Displacement

To assess basal contractility during hiPSC-EP network formation, 3D bead displacements were quantified in single cells and small multicellular clusters before and after cytochalasin-D treatment. Mean contractile displacement exhibited a non-linear exponential decay with increasing hydrogel shear storage modulus (Figure 2A–B), with smaller displacements observed in stiffer hydrogels. Multicellular clusters (2–4 cells) generated significantly greater total displacements than single cells, particularly in compliant hydrogels, although the per-cell contribution was reduced (Figure 2A,C; Table 1).

**Table 1.**
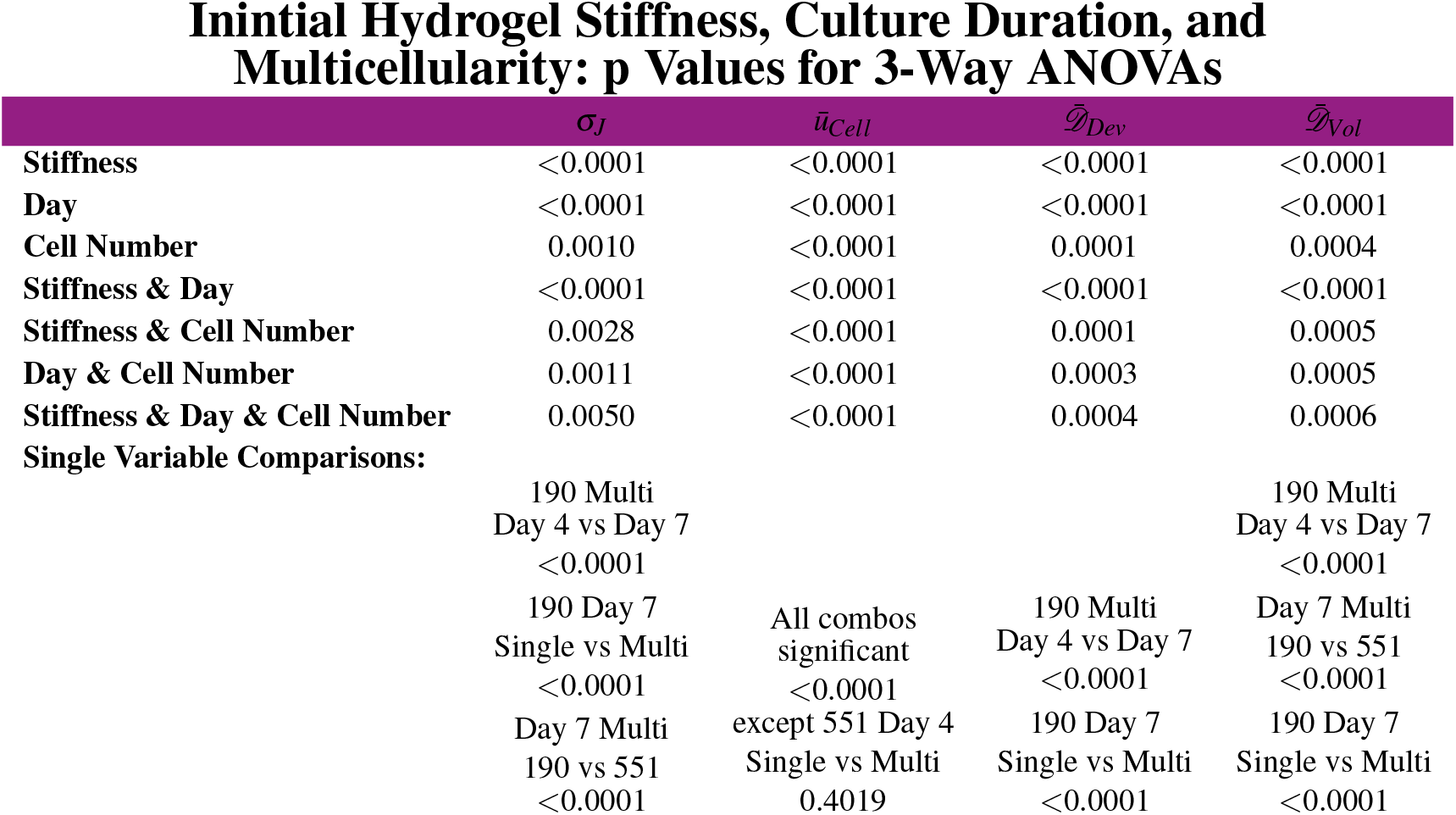
Culture duration, cell number, and hydrogel stiffness significantly influence hydrogel deformation and cellular contractility metrics. Three-way ANOVA with Tukey post-test evaluated effects of culture duration (Day 4 vs Day 7), cell number (Single vs Multi), and hydrogel stiffness (190 Pa vs 551 Pa) on hydrogel deformation, average cellular contractile displacement, deviatoric strain index, and volumetric strain index. Reported values include p values for each factor relative to the indicated dependent variable, followed by significant pairwise comparisons.

**Figure 2.**
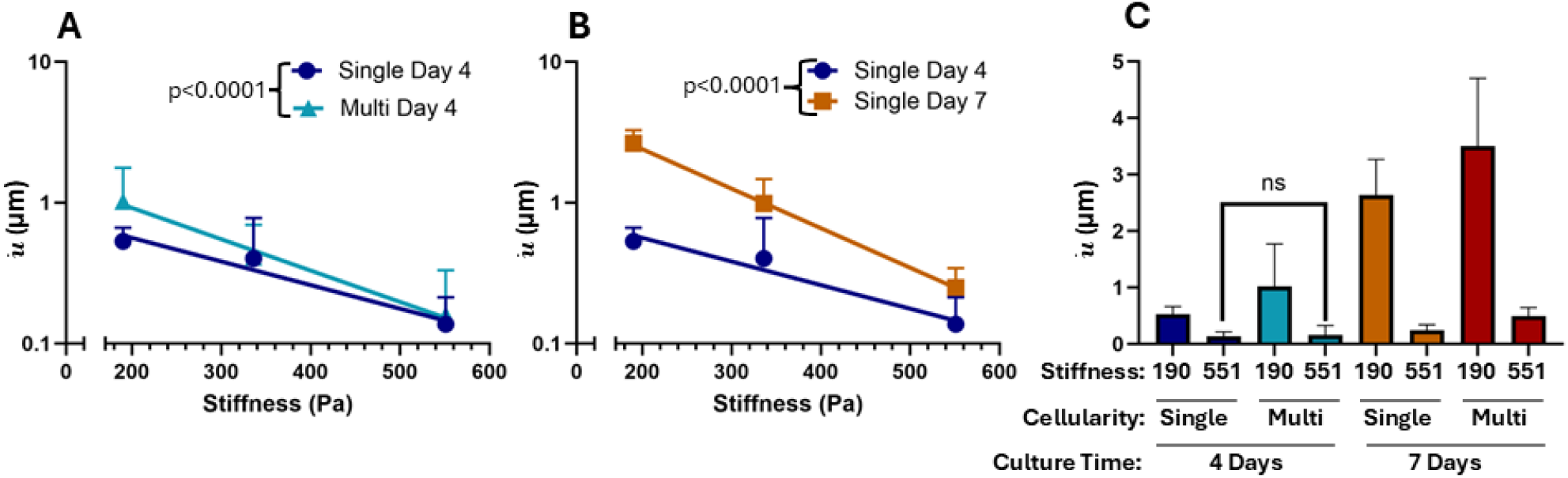
Hydrogel stiffness, culture duration, and cell number regulate cellular contractile displacement. A–B) Mean ± SEM displacement versus hydrogel stiffness with nonlinear regression (weighted one-phase decay, plateau = 0). Curve differences tested by sum-of-squares F test. A) Multi-cell Day 4: *d*(*G*) = 2.561*e*^−0.005125*G*^, R^2^ = 0.3381; Single-cell Day 4: *d*(*G*) = 1.218*e*^−0.003859*G*^, R^2^ = 0.3537; curves differ (p<0.0001). B) Single-cell Day 7: *d*(*G*) = 8.738*e*^−0.006462*G*^, R^2^ = 0.6412, significantly greater than Single-cell Day 4 (p<0.0001). C) Three-way ANOVA (190 and 551 Pa; Tukey post-test) showed significant effects of stiffness, culture duration, and cell number on basal contractile displacement (Table 1); all pairwise comparisons were significant (p<0.0001) except the comparison marked “ns.”

**Figure 3.**
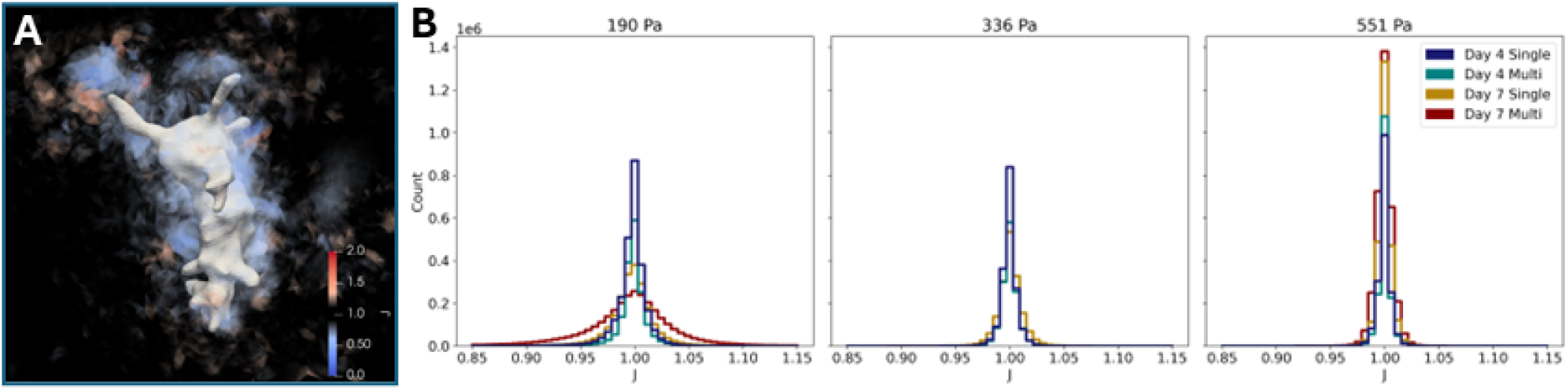
NorHA Hydrogels exhibited substantial spatial variations in fractional volume changes. A) Representative figure of a 190 Pa NorHA hydrogel containing a multi-cell group cultured for 7 days, showing spatial variations in local hydrogel *J* values, which ranged from less than 0.9 to more than 1.1. B) Histograms depicting the number of hydrogel finite elements with each *J* value ranging from *J*=0.85 to 1.15 per experimental group.

Extending culture duration from 4 to 7 days further increased basal contractility, most notably in compliant hydrogels (Figure 2B–C). This increase coincides with endothelial maturation and actin cytoskeletal development previously observed in hiPSC-EPs during this time frame.[9]

Collectively, hydrogel stiffness, cell number, and culture duration each contributed significantly—both independently and combinatorially—to basal contractile displacement (Figure 2C; Table 1). While stiffness and culture maturity exerted the largest effects, multicellularity had a smaller but significant influence, indicating that hiPSC-EP contractility is dynamically modulated by both intrinsic maturation and mechanical context.

### (b) Volumetric Variations During NorHA Hydrogel Deformation

EDP-crosslinked NorHA hydrogels are known to swell, indicating partial water mobility and material compressibility [30, 31]. Accordingly, we quantified local volumetric deformation during basal contraction by computing the Jacobian (*J*), where *J* = 1 represents incompressibility. Initial hydrogel stiffness significantly influenced volumetric deformation in a non-linear manner (Figure (b); Table 1). Because stiffness was modulated *via* crosslinker concentration—which also governs porosity—stiffness was implemented as the representative material parameter throughout this study.[30–32]

Notably, in the most compliant (190 Pa) hydrogels, 20% of elements exhibited substantial volume changes (*J* ∉ [0.9, 1.1]; Figure (b)), demonstrating pronounced cell-induced compressibility. Across all conditions, variations in *J* were primarily driven by initial hydrogel stiffness and culture time, with a smaller but significant contribution from multicellularity (Table 1), indicating that hiPSC-EPs actively remodel their hydrogel microenvironment through contractility-mediated and time-dependent ECM remodeling mechanisms.

### (c) Kinematic Analyses of NorHA Hydrogels During Basal Contraction

Because hydrogels exhibited significant compressibility, strain was decomposed into deviatoric and volumetric components. Both showed non-linear dependence on hydrogel stiffness, with substantially higher values in more compliant hydrogels across all conditions (Figure 4A–B,D–E). Lower stiffness produced multi-fold increases in hiPSC-EP–induced strain, reflecting greater deformation per unit force in compliant matrices.

**Figure 4.**
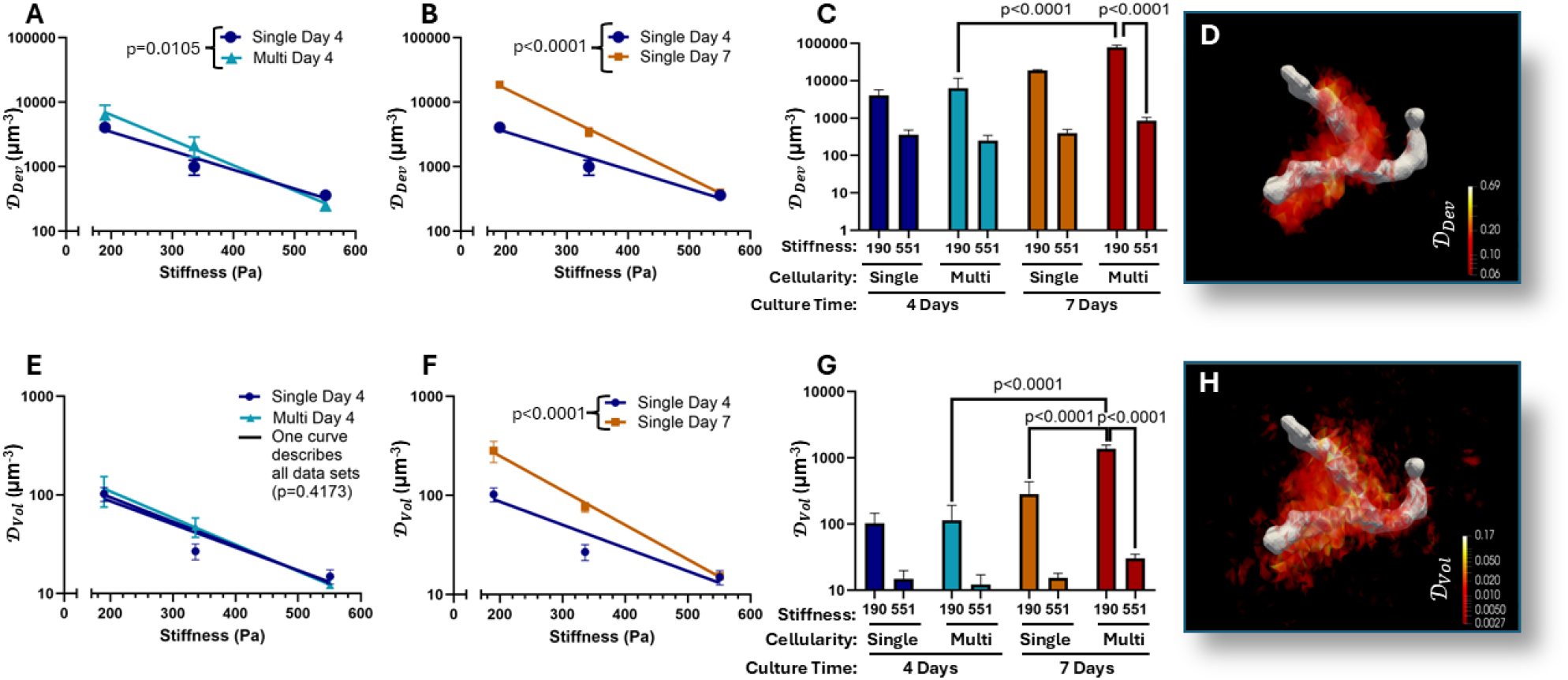
Hydrogel stiffness, culture duration, and cell number regulate deviatoric and volumetric strain indices. A–B, E–F) Mean ± SEM strain index versus hydrogel stiffness with nonlinear regression (weighted one-phase decay, plateau = 0); curve differences tested by sum-of-squares F test. C, G) Three-way ANOVA with Tukey post-test for 190 and 551 Pa hydrogels (Table 1); significant comparisons indicated by brackets. A) Multi-cell Day 4 groups showed greater deviatoric strain in compliant hydrogels (*σ* (*G*) = 38685*e*^−0.009019*G*^, R^2^ = 0.6312) than Single-cell Day 4 groups (*σ* (*G*) = 13381*e*^−0.006758*G*^, R^2^ = 0.7431; p=0.0105). B) Single-cell Day 7 groups exhibited greater deviatoric strain (*σ* (*G*) = 134931*e*^−0.01062*G*^, R^2^ = 0.9178) than Single-cell Day 4 groups (p<0.0001). C) Stiffness, culture duration, and cell number significantly contributed to deviatoric strain index. D) The largest 90th percentile of deviatoric strain localized near hiPSC-EPs. E) Single- and Multi-cell Day 4 volumetric strain were described by a single curve (*σ* (*G*) = 297.2*e*^−0.005707*G*^, R^2^ = 0.6632; p=0.4173). F) Single-cell Day 7 groups showed greater volumetric strain (*σ* (*G*) = 1245*e*^−0.008017*G*^, R^2^ = 0.8075) than Single-cell Day 4 groups (*σ* (*G*) = 254.0*e*^−0.005390*G*^, R^2^ = 0.6819; p<0.0001). G) Stiffness, culture duration, and cell number significantly contributed to volumetric strain index. H) The largest 90th percentile of volumetric strain also localized near hiPSC-EPs.

Deviatoric strain was higher in multicellular groups than single cells, particularly in compliant hydrogels, with synergistic effects emerging by day 7 (Figure 4A,C; Table 1). Three-way ANOVA confirmed that hydrogel stiffness, cell number, and culture duration independently and combinatorially contributed to deviatoric strain (Figure 4C; Table 1), indicating that hiPSC-EPs impose shape-changing strains that evolve with mechanical context and maturation

Volumetric strain showed no multicellular effect at day 4, collapsing onto a single stiffness-dependent curve (Figure 4D). By day 7, multicellularity significantly increased volumetric strain in the most compliant (190 Pa) hydrogels, with effects exceeding 11-fold relative to single cells (Figure 4F; Table 1). Culture duration further amplified volumetric strain, particularly in compliant hydrogels, yielding >12-fold increases from day 4 to 7 (Figure 4E–F).

Although deviatoric strain dominated—averaging 40× volumetric strain—the presence of measurable volumetric strain and Jacobian variations confirms that compressibility is non-negligible. These results demonstrate that both strain components must be considered to accurately characterize hiPSC-EP–mediated deformation in crosslinked NorHA hydrogels.

### (d) Spatial Variation in Hydrogel Stiffness

By implementing our inverse modeling approach which includes terms for deviatoric and volumetric strain in the forward solve [19], the modulus of each cellular system embedded in the softest, 190 Pa, hydrogels was found to vary spatially (Figure 5). As extensively described in the above methods section, only the 190 Pa hydrogels could be analyzed due to low signal/noise ratio for displacements in the stiffer hydrogels. In the 190 Pa hydrogels, as culture time increased, the amount of stiffening significantly increased, which suggested that the hiPSC-EPs had a longer time to secrete ECM macromolecules such as laminin and collagen into the hydrogel [33]. Concurrently, multicellularity and culture time both significantly decreased the amount of degradation that was imparted on the hydrogels, suggesting that iPSC-EPs decrease the secretion of MMPs [5, 14–16]. Moreover, modulus was significantly higher close to the cellular surfaces, with day 7 multi-cell samples stabilizing into a linear trend for the relationship of distance from cell vs. *log*_2_ fold change of modulus. These results suggest that the iPSC-EPs cause the hydrogels to become stiffer by both increasing stiffening and decreasing degradation over time.

**Figure 5.**
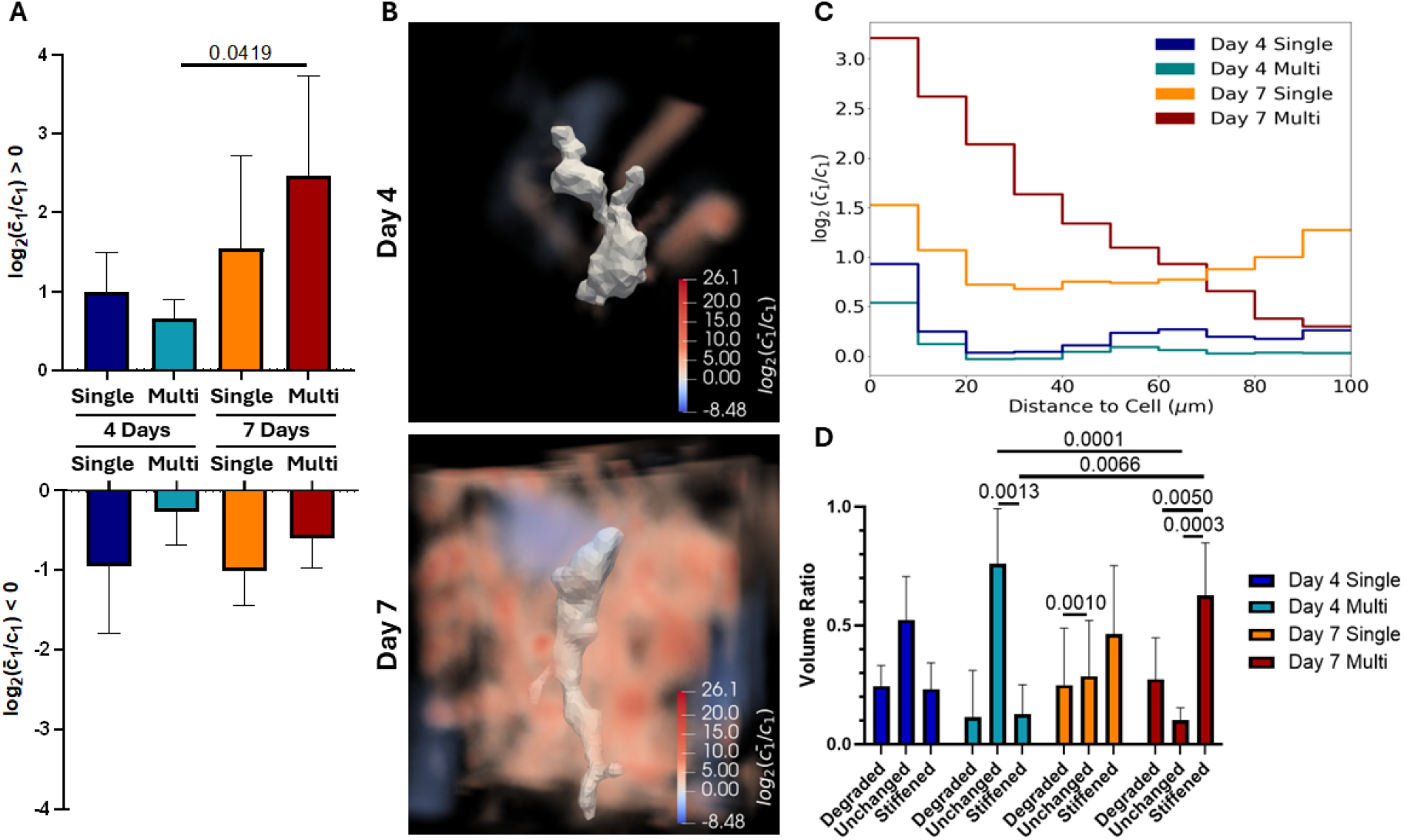
Hydrogel stiffness varies with culture time, multicellularity, and distance from cellular surfaces. A) Mean *log*_2_ fold change in modulus ± SD for stiffened (*>* 0) and degraded (< 0) regions. Two-way ANOVA showed a significant effect of culture time in stiffened regions (p=0.0156; Tukey post-test indicated pairwise differences). In degraded regions, the interaction between multicellularity and culture time was significant (p=0.0367). B) Representative images illustrating increased pericellular stiffening in multicellular hiPSC-EP groups from day 4 to day 7. C) In day 7 multicellular cultures, stiffness decreased with distance from cell surfaces (*y* = −0.03130*x* + 2.994, *r*^2^ = 0.9529). Three-way ANOVA identified significant effects of culture time, multicellularity, and distance individually (*p* < 0.0001) and in combination (*p* < 0.0001). D) Mean volume ratio ± SD of degraded (< global third quartile), stiffened (> first quartile), and unchanged regions. Three-way ANOVA showed significant variation among modulus groups (*p* = 0.0079), with strong effects of culture time (*p* < 0.0001) and multicellularity (*p* = 0.0171); Tukey post-test indicated pairwise differences.

### (e) Strain Energy Density

Since modulus is high near the cellular surfaces and basal contractility displacements of the hydrogel are also more prominent near the cellular surfaces, it is not surprising to find that strain energy density is significantly higher in zones near cellular surfaces (Figure 6). Furthermore, culture time and multicellularity both significantly increase strain energy density, reflecting the progressive maturation and cooperative contractility of hiPSC-EPs that leads to larger deformations and higher elastic energy in the surrounding matrix.

**Figure 6.**
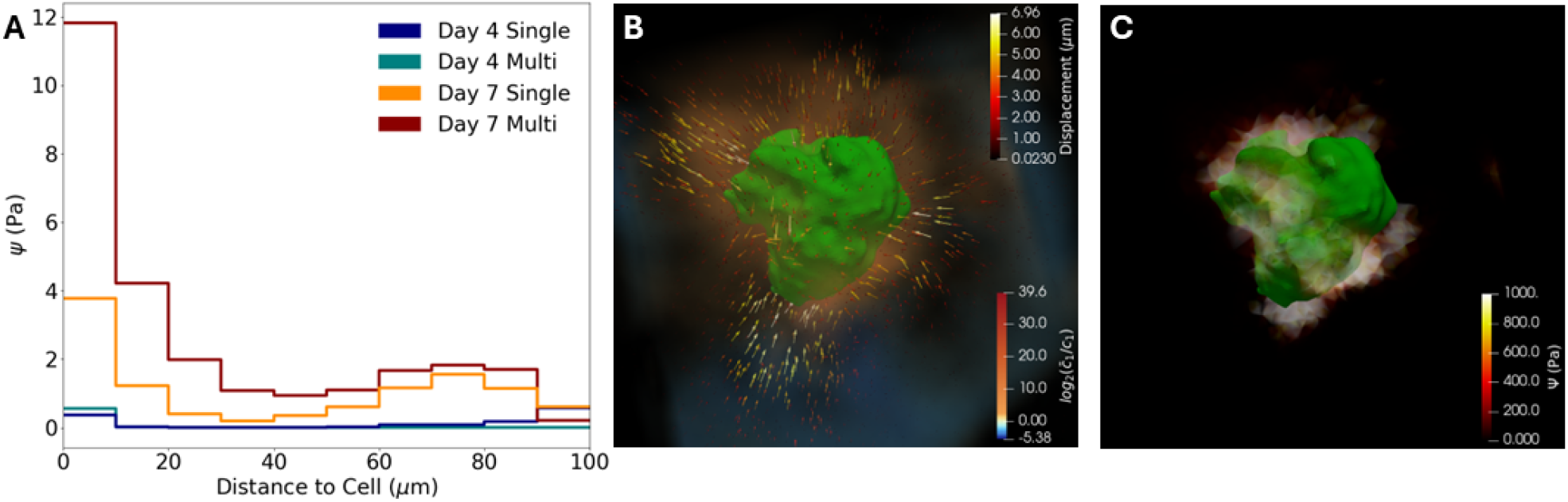
Strain energy density is highest near cellular surfaces where contractile displacements and hydrogel modifications are greatest. A) Step plot showing strain energy density as a function of distance from the cell. A 3-way ANOVA found culture time, multicellularity, and distance from the cell to be significant contributors both individually and in combination (*p* < 0.0001). B) Representative Day 7 multicellular cluster showing hydrogel contractile displacement (*µ*m) and modulus modification 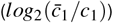. C) The same system illustrating that regions of highest strain energy occur near cellular surfaces where ECM modification and contractility are greatest.

### (f) Traction Forces

As seen with strain energy density, the total traction force per cell system was significantly increased by longer culture time and multicellularity (Figure 7), suggesting that as the culture develops, the hiPSC-EPs’ actomyosin networks mature and exert stronger contractile forces on the surrounding matrix [34], which is further supported by our previous findings that actin network formation in these cells develops over time. [9] In addition, within the day 7 multicellular samples, punctae of high traction force magnitudes could be seen, while these punctae were not present in single celled samples. This suggests that this method may be able to capture in 3D the locations of focal adhesion hot spots that develop during cell migration [35] which we have previously seen in fixed staining of viculin in these cells. [9]

**Figure 7.**
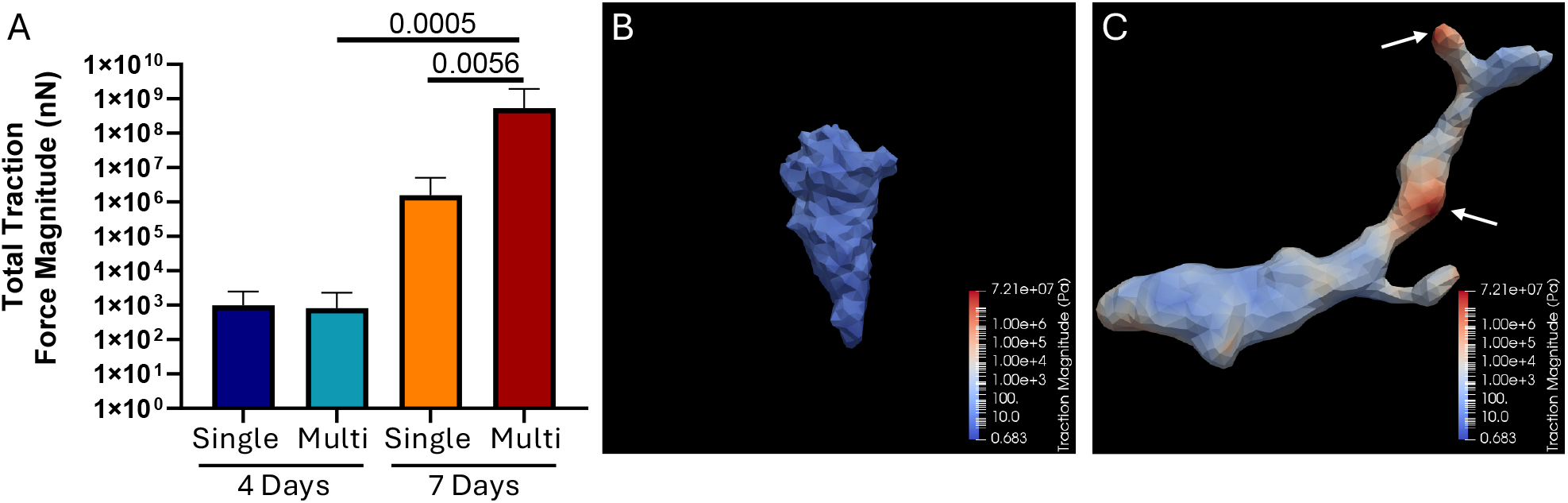
Traction forces increase with culture time and multicellularity. A) Mean ± standard deviation of total traction force magnitude (nN) for each cell system. A two-way ANOVA on *log*_10_-transformed data found significant effects of culture time (*p* = 0.0003), multicellularity (*p* = 0.0199), and their interaction (*p* = 0.0195). B) Representative Day 7 single-cell image showing uniformly low traction forces. C) Representative Day 7 multicellular cluster showing localized high-traction hotspots at the cell surface (white arrows).

## 4. Discussion and Conclusions

### (a) Findings

Our long-term goal is to understand how hiPSC-EPs modify their surroundings as they assemble into lumenized networks in 3D crosslinked NorHA hydrogels, ultimately enabling the design of angiogenic biomatrices that promote robust microvascularization in hiPSC-derived constructs. As an initial step, this study examined the transition from single cells to small multicellular clusters during the early days after hiPSC-EP embedding. Hydrogel stiffness, culture duration, and cell number significantly influenced basal contractile displacement magnitude and were also major determinants of ECM remodeling, hydrogel strain energy density, and traction force magnitude at cell surfaces.

A key finding is that hiPSC-EPs substantially remodel crosslinked NorHA hydrogels. Cells are known to modify hydrogels through ECM deposition, enzymatic degradation, and contractile forces [5, 14–16], altering local or bulk mechanical properties across many systems [14–16, 18, 36]. However, detailed three-dimensional micron-scale measurements of scalar deformation in NorHA have been lacking. Most prior models treat NorHA as incompressible or nearly incompressible [37–39] despite documented swelling behavior [30, 31].

Previous 3D-TFM studies of endothelial sprouting accounted for hydrogel compressibility when estimating traction forces but did not incorporate spatial heterogeneity in the cell-modified hydrogel modulus [40, 41]. More broadly, although compressibility has been addressed through deviatoric invariants in several 3D strain-mapping studies, spatial variation in hydrogel modulus has rarely been incorporated into traction or strain energy calculations [42–45].

Here, building on techniques we previously developed [10, 19], full 3D scalar deformation was quantified in crosslinked NorHA hydrogels and applied to resolve micron-scale spatial heterogeneity in hydrogel modulus arising from cellular remodeling. These local changes are consistent with prior observations that cells deposit ECM proteins such as collagen and laminin with mechanical properties distinct from the native hydrogel [5, 14–16]. This spatially varying modulus was then incorporated into strain energy density and traction force calculations. To our knowledge, this is the first study linking initial hydrogel stiffness, culture duration, and multicellularity to cell-mediated micron-scale heterogeneity in modulus and incorporating that heterogeneity to more accurately estimate strain energy and traction forces in NorHA hydrogels.

A central hypothesis was that multicellularity would amplify the effects of hydrogel stiffness and culture duration on hiPSC-EP contractility and hydrogel remodeling. In day 4 cultures, multicellularity increased most parameters but did not produce clear synergistic effects. By day 7, however, synergism emerged in all strain energy and traction force measurements. Although clusters contained only 2–4 cells, these results suggest that even small groups can drive collective matrix and cytoskeletal remodeling over time, producing stiffer matrices, more robust cytoskeletal structures, and the larger strain energies and traction forces observed [35].

### (b) Study Limitations

The present investigation examined the transition of hiPSC-EPs from the fully relaxed CytoD-treated steady state to the basal steady state to determine basal tonus. This scope was limited by the time required to acquire high-resolution full-3D images of the cells and hydrogels. The data show that the more compliant crosslinked NorHA hydrogels became increasingly compressible over the culture period of embedded hiPSC-EPs. Because minute-scale measurements during cell relaxation were not possible, time-dependent hydrogel models could not be explored. Previous work has assumed that EDP-crosslinked NorHA hydrogels behave as incompressible neo-Hookean materials under steady-state conditions, [37] consistent with many other hydrogel systems. [46, 47] The porosity of cell-free NorHA hydrogels is well characterized, [32] and these hydrogels exhibit poroelastic behavior over relevant time scales. [48–50] Thus, water transport through the porous network may be the primary mechanism underlying the compressibility observed during hiPSC-EP basal contraction [51–53] Recent advances in light-sheet confocal microscopy have greatly increased the speed of high-precision full-3D imaging, [54] which may enable future investigation of time-dependent hydrogel dynamics in cellularized systems.

### (c) Conclusions

We demonstrate that key environmental conditions strongly influence hiPSC-EP–mediated microenvironmental remodeling of hydrogels. Both volumetric and deviatoric strain depend on 3D culture parameters, producing synergistic changes in system strain energy density. Over time, matrix remodeling locally stiffens the hydrogel, generating regions of elevated stiffness near cells. The combination of increased stiffness and elevated strain energy produces large traction forces in long-culture, multicellular systems, with localized regions of even greater force. Further quantification of these effects will deepen understanding of the biomechanical mechanisms governing hiPSC-EP self-assembly into interconnected lumenized networks and support the design of angiogenic biomaterials that promote vascularization in hiPSC-derived tissue constructs.

## Ethics

Experiments were carried out in accordance with the University of Texas at Austin regulations and NIH guidelines.

## Data Accessibility

Data that support the findings of this study are available at the Texas Data Repository, https://doi.org/10.18738/T8/65QOHL.

## Authors’ Contributions

TMW: lead — data curation, formal analysis, software, validation, visualization, writing/original draft; equal — funding acquisition, investigation, methodology; supporting — project administration. JH: equal — investigation, methodology; supporting — data curation. GP: supporting — formal analysis, software, visualization, writing/review & editing. JZ: lead — resources; equal — conceptualization, funding acquisition, methodology, project administration, supervision, writing/review & editing; supporting — formal analysis. MS: lead — project administration, resources, supervision; equal — conceptualization, funding acquisition, methodology, writing/review & editing; supporting — formal analysis.

## Acknowledgements

Imaging occurred at the Center for Biomedical Research Support Microscopy and Flow Cytometry Facility at the University of Texas at Austin (RRID: SCR_021756).

